# Precise Slow Oscillation-Spindle Coupling Promotes Memory Consolidation in Younger and Older Adults

**DOI:** 10.1101/268474

**Authors:** Beate E. Muehlroth, Myriam C. Sander, Yana Fandakova, Thomas H. Grandy, Björn Rasch, Yee Lee Shing, Markus Werkle-Bergner

## Abstract

Memory consolidation during sleep relies on the precisely timed interaction of rhythmic neural events. Here, we investigate differences in slow oscillations (SO) and sleep spindles (SP) and their coupling across the adult human lifespan and ask whether observed alterations relate to the ability to retain associative memories across sleep. We demonstrate that the fine-tuned SO–SP coupling that is present in younger adults diffuses with advanced age and shifts both in time and frequency. Crucially, we show that the tight precision of SO–SP coupling promotes memory consolidation in younger and older adults, and that brain integrity in source regions for the generation of SOs and SPs reinforces this beneficial SO–SP coupling in old age. Our results reveal age-related differences in SO–SP coupling in healthy elderly individuals. Furthermore, they broaden our understanding of the conditions and the functional significance of SO–SP coupling across the entire adult lifespan.

## Introduction

Research in animals and younger adults yields converging evidence that newly acquired information is reprocessed during deep non-rapid eye movement (NREM) sleep. According to the framework of system consolidation^1,2^ declarative memories are initially processed by a fast learning system in the hippocampus that binds information into a transient representation^3^. Only repeated reactivation of memories in the hippocampus then eventually allows transfer to a long-term store in the neocortex^4–6^. But how does the brain ensure precise communication between cortical modules that allows for the reorganization of memory traces during sleep?

During sleep, effective communication is enabled by the fine-tuned hierarchical succession of several brain rhythms governed by large-amplitude, low-frequency oscillations, so-called slow oscillations (SO; 0.5–1 Hz) that mainly emerge from prefrontal brain areas^7–10^. Via cortico-thalamic projections, neurons in the thalamic reticular nucleus are activated during the depolarized up-states of SOs resulting in the initiation of fast sleep spindles (SP; 12.5–16 Hz)^11–14^. These fast SPs, in turn, synchronize cortical activity in a precise spatiotemporal manner. As the excited neurons exhibit a massive influx of Ca^2^+, synaptic changes become facilitated^15^. Further nesting of hippocampal high–frequency bursts, known as sharp-wave ripples, within the excitable troughs of fast SPs^16–18^ optimizes the hippocampo–neocortical dialogue. A second SP type, so-called slow SPs (9–12.5 Hz), occurs during a time window in which neural depolarization (i.e., the SO up-state) transitions into hyperpolarized down-states^12,19,20^. A specific role of slow SPs in the consolidation of memories is not yet established^2,19,21^.

Typically, SOs and SPs are studied with regard to their independent contribution to consolidation mechanisms. However, as system consolidation is assumed to rely on a tight coupling of SOs, SPs, and sharp-wave ripples^4,22^, recent studies have started to focus on the exact timing of brain rhythms^11,19,20,23,24^ and its functional significance for sleep-dependent memory consolidation^19,24–27^. Recent experiments in rats demonstrated that reinforcing the coordinated pattern of SOs, SPs, and sharp-wave ripples boosts memory consolidation of object locations^25^, whereas suppression of these patterns has disruptive effects on sleep-dependent memory consolidation^26^. In humans, SO–SP coupling is enhanced by prior learning^19,24^ and is triggered by replaying previously learned material during sleep^28^. Although first studies have highlighted the involvement of SO–SP coupling in memory consolidation, particularly in humans, evidence for this link is still sparse. Further, it remains to be established under which circumstances the coordination of SOs and SPs is most beneficial in supporting the sleep-dependent reorganization and stabilization of memory contents.

SO–SP coupling, for instance, can be critically impaired in disrupted sleep. This is particularly common in elderly adults who often lack the deepest NREM sleep stage, so-called slow-wave sleep^29–31^. In old age, SOs and SPs themselves appear less often and with reduced amplitudes^32–34^. Age-related changes in brain structure possibly contribute to the observed sleep alterations by impairing SO and SP generation and propagation^33,35–38^. This may also impact SO–SP coupling itself, thereby resulting in impaired memory stabilization during sleep^39^. Indeed, several studies report age-related losses in overnight memory retention^40–42^, which can be linked to reduced slow-wave sleep^35,36^ and a decline in SP occurrence^43^. In light of the growing interest in the interaction of SOs and SPs, very recent evidence highlights that SO–SP coupling is indeed dispersed in elderly adults, resulting in impaired overnight memory consolidation^39^. As this disturbed nesting of SOs and SPs may constitute a possible target for clinical interventions^44^, the preconditions and the functional significance of SO–SP coupling across the entire adult lifespan require investigation.

Following this rationale, it was our aim to examine age-related changes in the coupling of SOs and both slow and fast SPs. We asked whether observed alterations can be explained by structural brain atrophy in source regions of SO and SP generation and whether the resulting SO–SP dispersion negatively impacts the ability to retain associative memories across sleep. In the present study, younger and older adults completed an associative memory paradigm, consisting of a learning session of scene–word pairs on the first day and a delayed cued-recall task on the following day (Figure 1). During the nights before and after learning, sleep was monitored using ambulatory polysomnography (PSG). Structural brain integrity was assessed by voxel-based morphometry (VBM) of structural magnetic resonance images (MRI). By comparing the PSG recordings of younger and older adults, we identify age-related differences in the coordination of SOs and SPs. Using estimates of gray matter volume we further establish the role of structural brain integrity in the coordination of SOs and SPs. Finally, we relate the pattern of SO–SP coupling to the behavioral measure of overnight memory retention and show that the observed ‘aged’ SO–SP coordination indeed explains deficient memory consolidation in old age.

**Figure 1.**
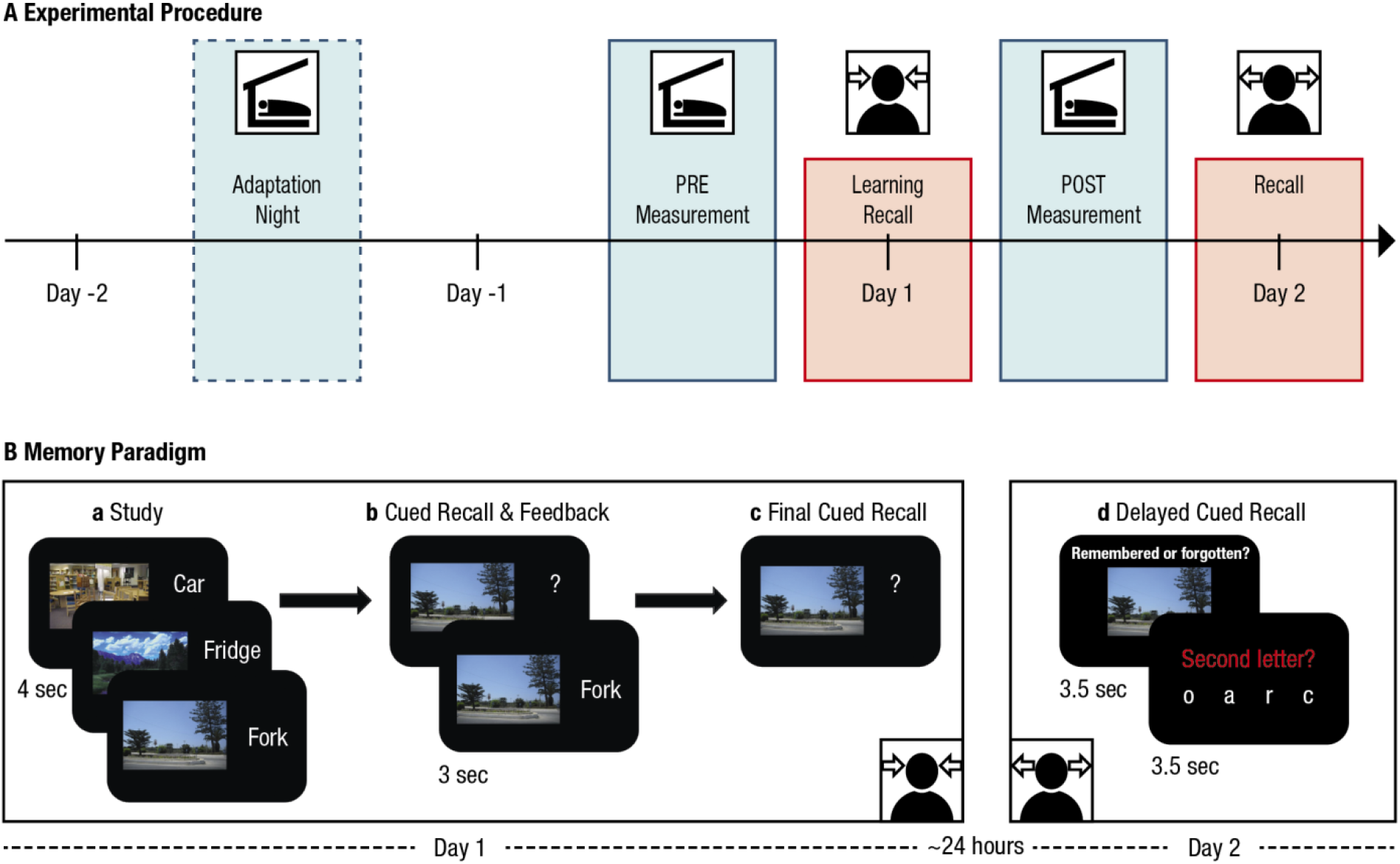
A. Experimental procedure. The memory task at the core of the experiment consisted of a learning phase and an immediate recall on Day 1 as well as a delayed recall approximately 24 hours later (red boxes). Sleep was monitored in the nights before (PRE) and after (POST) learning using ambulatory polysomnography (blue boxes). A prior adaptation night (dashed blue box) familiarized the participants with the sleep recordings. B. Memory paradigm (cf. Fandakova et al., 2018^63^). (a) During study, participants were instructed to remember 440 (younger adults) or 280 scene–word pairs (older adults). (b) During the cued-recall and feedback phase the scene was presented as a cue to recall the corresponding word. Irrespective of recall accuracy, the original pair was presented again to allow for re-study. The whole cued-recall and feedback cycle was performed once in younger and twice in older adults. (c) During final recall, scenes again served as cues to recall the corresponding word, but no feedback was provided. (d) Delayed cued-recall took place approximately 24 hours later. Participants were presented with the scenes only and had to indicate if they still remembered the associated word. Afterwards they had to select the corresponding second letter of the word to verify their true memory of the associate.

## Results

### Rhythmic neural events during NREM sleep are affected by aging

Sleep architecture changes drastically across the adult lifespan^30^. To capture these changes, we recorded electrical neuronal activity during sleep before and after learning scene–word pairs using ambulatory PSG devices at participants’ homes (Methods). In the following, we focus on sleep after learning only. In comparison to younger adults, we found a lower proportion of slow-wave sleep in older adults, accompanied by a higher proportion of lighter NREM sleep stages 1 and 2 (Table 1). As almost one third of our older sample (*n* = 9) failed to meet the amplitude criteria of slow-wave sleep when visually scoring the PSG data^45^, we combined NREM sleep stage 2 and slow-wave sleep for further analyses.

**Table 1.** Age differences in sleep architecture

|  | Younger Adults – Median<br>[1st quartile; 3rd quartile] | Older Adults – Median<br>[1st quartile; 3rd quartile] | *z* | *p* |
| --- | --- | --- | --- | --- |
| TST (min) | 457.00 [420.0; 508.5] | 417.50 [370.75; 447.25] | 2.60 | .009 |
| WASO (min) | 3.43 [0.73; 5.66] | 9.09 [4.96; 15.77] | –3.81 | <.001 |
| Stage 1 (%) | 4.31 [3.07; 5.88] | 7.57 [4.72; 10.20] | –3.19 | .001 |
| Stage 2 (%) | 50.69 [44.96; 56.92] | 63.99 [61.09; 69.96] | –5.45 | <.001 |
| SWS (%) | 20.56 [16.62; 26.61] | 3.80 [0.00; 12.67] | 5.38 | <.001 |
| REM (%) | 24.49 [21.6; 26.69] | 21.10 [17.01; 24.53] | 2.29 | .022 |

Note. TST: total sleep time; WASO: wake after sleep onset; SWS: slow-wave sleep; REM: rapid eye-movement sleep.

Within NREM sleep we were particularly interested in the occurrence of SOs, as well as slow and fast SPs (slow SP: 9–12.5 Hz, fast SP: 12.5–16 Hz, cf. Methods). In line with the previously reported topography of slow and fast SPs^20^, we focused our analyses on frontal slow SPs and central fast SPs. Here, we observed the expected reduction in the occurrence of frontal SOs and slow SPs (SO number: *z* = 2.6, *p* = .009, *Md*_YA_ = 1140.5, *Md*_OA_ = 970.50; slow SP number: *z* = 3.41, *p* < .001, *Md*_YA_ = 275.75, *Md*_OA_ = 141.50; slow SP density: *z* = 3.16, *p* = .002, *Md*_YA_ = 0.83 events/min, *Md*_OA_ = 0.48 events/min) as well as central fast SPs (fast SP number: *z* = 5.89, *p* < .001, *Md*_YA_ = 511, *Md*_OA_ = 162; fast SP density: *z* = 5.32, *p* < .001, *Md*_YA_ = 1.70 events/min, *Md*_OA_ = 0.58 events/min) (see also Supplementary Table 1 and Supplementary Figure 1). When accounting for the time spent in NREM sleep, the density of SOs (events/min), however, was not significantly reduced in older compared to younger adults (SO density: *z* = 1.53, *p* = .127, *Md*_YA_ = 3.72 events/min, *Md*_OA_ = 3.36 events/min).

### Slow and fast SPs sustain their dependency on SOs in older adults

To determine whether SP events indeed coincide with the detected SOs, we calculated the amount of SPs whose center occurred in an interval of ± 1.2 s around the trough of the identified SOs (Figure 2A). The time window was chosen to cover one whole SO cycle (0.5–1 Hz, i.e., 1–2 s). To cross-check, we also calculated the proportion of SOs with SPs whose center occurred within ± 1.2 s around the respective trough of the oscillation (Figure 2B). In both age groups we identified a comparable percentage of slow SPs coupled to SOs *(t(48)* = −0.67, *p* = .508, M_YA_ = 57.13 %, *M*_OA_ = 59.60 %, Figure 2A) and, vice versa, a similar amount of SOs co-occurring with slow SPs (t(50) = 1.61, *p* = .115, *M*_YA_ = 14.36.75 %, *M*_oa_ = 11.18 %, Figure 2B). Although we found a reduced proportion of SOs occurring in coordination with fast SPs in older adults (t(48) = 4.06, *p* < .001, *M*_YA_ = 14.43 %, *M*_OA_ = 7.34 %), the occurring fast SPs were as likely to coincide with SOs in older as in younger adults (t(52) = –2.08, *p* = .043, not significant after applying Bonferroni correction for multiple comparisons, *M*_YA_ = 31.92 %, *M*_oa_ = 37.92 %). This suggests a generally constant pattern of SP–SO coupling across adulthood, with potentially the same mechanisms underlying SP generation. SOs might partly forfeit their ability to initiate fast SP generation in old age. However, those SPs that do occur in older adults are still bound to the occurrence of SOs.

**Figure 2.**
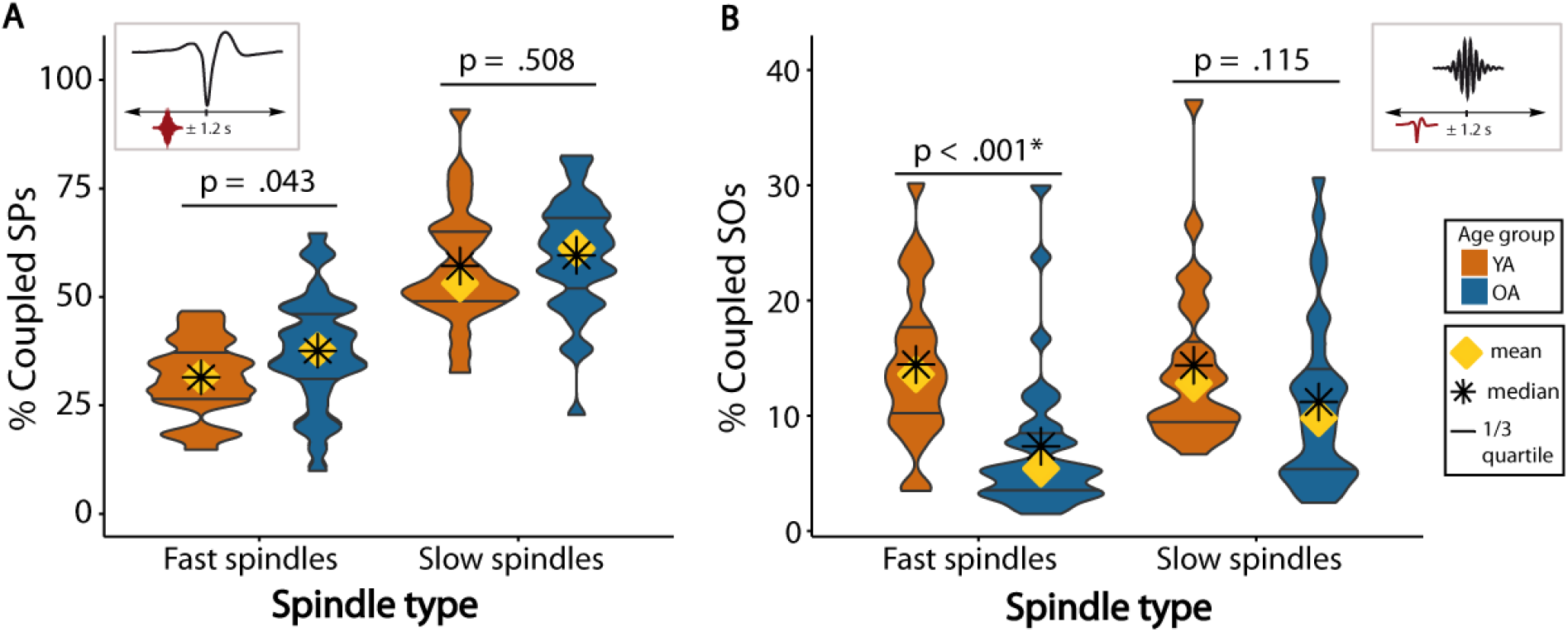
Fast and slow SPs coincide with SOs in both younger and older adults. Asterisks mark significance after correction for multiple comparisons. (A) About one third of all fast SPs and more than half of all slow SPs occur within an interval of ± 1.2 s around the trough of a SO. (B) Only a small percentage of all appearing SOs is coupled to the occurrence of SPs. SOs in older adults (blue) occur less frequently within an interval of ± 1.2 s around fast SP centers than in younger adults. YA: younger adults; OA: older adults; SO: slow oscillation; SP: spindle.

Having established the general presence of SO–SP coordination in old age, we focused on the precise temporal association between SOs and SPs in the next step. We thus calculated peri-event time histograms (PETHs) of SOs co-occurring with fast and slow SP events (Figure 3, Supplementary Figures 3 and 4, cf. Methods). In short, these PETHs summarize the probability of fast and slow SP events occurring at a specific time point within ± 1.2 s around the SO trough. Stability of these estimates was tested by contrasting the PETHs with their randomly shuffled surrogates (cf. Methods). In younger adults, we demonstrate the expected preferential occurrence of fast SPs during the positive peaks preceding and following the SO trough (both clusters: *p* ≤ .012), whereas slow SPs were reliably enhanced during the up-to down-state transitions (both clusters: *p* ≤ .008, Figure 3A and 3C). Whereas in older adults neither fast nor slow SPs increased before the SO trough, SP increases following the trough resembled those of younger adults: slow SPs peaked during the up-to down-state transition (cluster *p* = .002) and fast SPs during the up-state (cluster *p* = .002, Figure 3B and 3D). While the fast SP increase in younger adults co-occurred with the positive SO peak (SO peak at 480 ms, significant positive cluster from 200 to 700 ms), fast SPs peaked before the SO peak in older adults (SO peak at 618 ms, significant positive cluster from 200 to 400 ms) and their temporal specificity was overall less pronounced.

**Figure 3.**
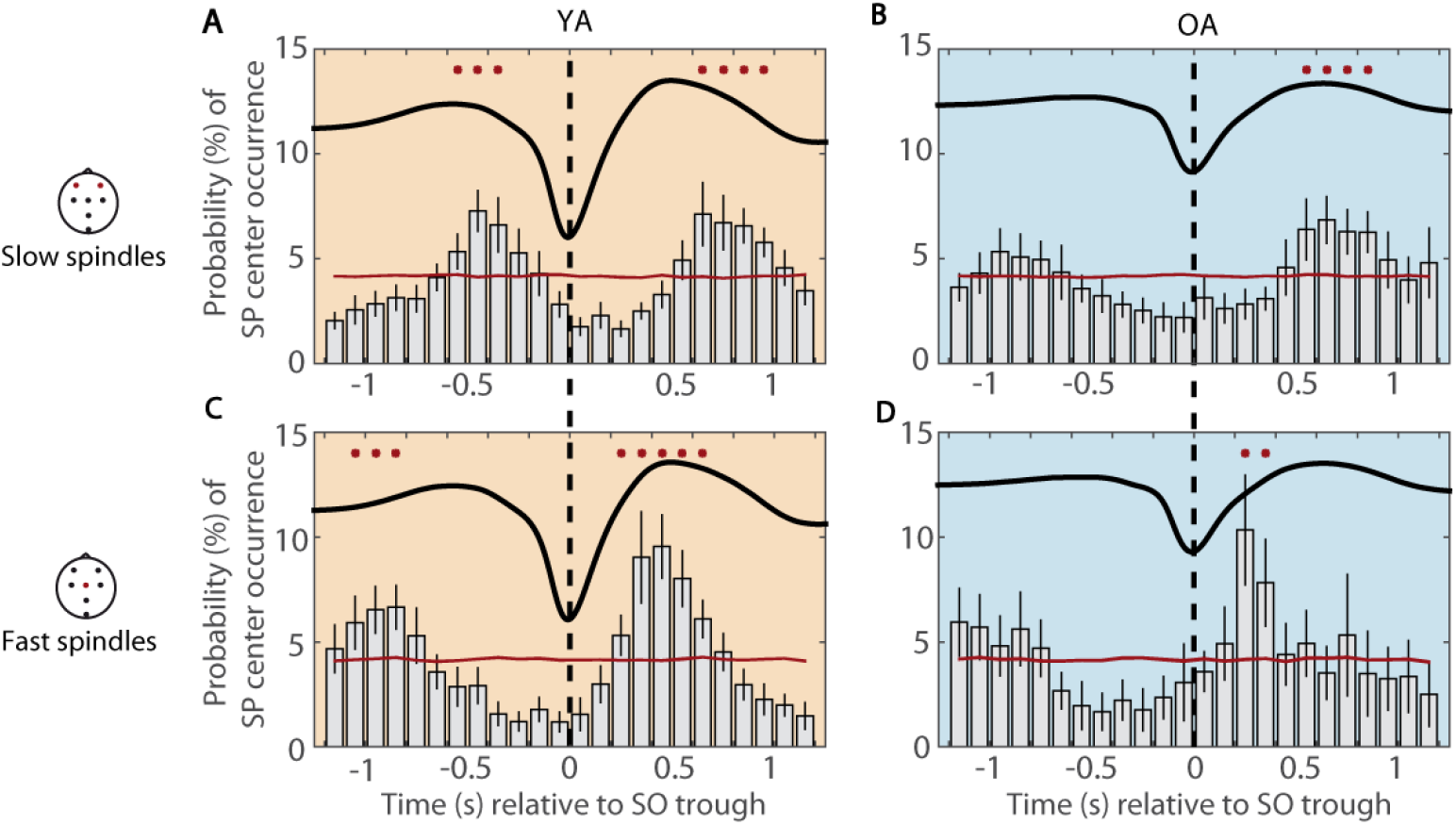
Temporal specificity of fast and slow SP occurrence. Peri-event time histograms of fast and slow SPs co-occurring with frontal SOs are depicted for both age groups. Standard errors of the 100-ms bins are included as black vertical lines. Reference distribution obtained after randomization of the data is shown in red. Red asterisks indicate significantly increased SP occurrence contrasted with the reference distribution (cluster-based permutation test, cluster α < 0.05, positive clusters only). Vertical dashed lines mark the SO trough. Average SOs are shown for each age group. (A) Frontal slow SPs in younger adults globally peak at the up-to down-state transition. (B) The frontal slow SP peak at the up- to down-state transition preceding the SO trough observed in younger adults disappears in older adults. (C) In younger adults, fast SPs prominently peak during the SO peak (SO peak at 480 ms, significant positive cluster: 200–700 ms). (D) In older adults, fast SPs are similarly modulated during the SO up-state as observed in younger adults but their occurrence is maximized before the SO has peaked (SO peak at 618 ms, significant positive cluster: 200–400 ms). YA: younger adults; OA: older adults; SO: slow oscillation; SP: spindle.

### SO–SP coupling differentially changes for fast and slow SPs in younger and older adults

Having established the overall maintenance of a time-specific SO–SP coordination in older adults, we further probed this pattern by comparing the oscillatory power in the SP frequency range for time segments with and without SOs (cf. Methods). During SO trials (centered on the trough of the respective SO ± 1.2 s), power in the SP frequency range indeed differed significantly from randomly selected intervals without SOs (for all clusters: *p* < .001, Figure 4B, Supplementary Figure 2). This effect was present in both age groups and suggests a coupling of SPs to specific phases of the underlying SO. Nevertheless, each age group displayed a distinct modulation of the effect (Figure 4B, Supplementary Figure 2).

**Figure 4.**
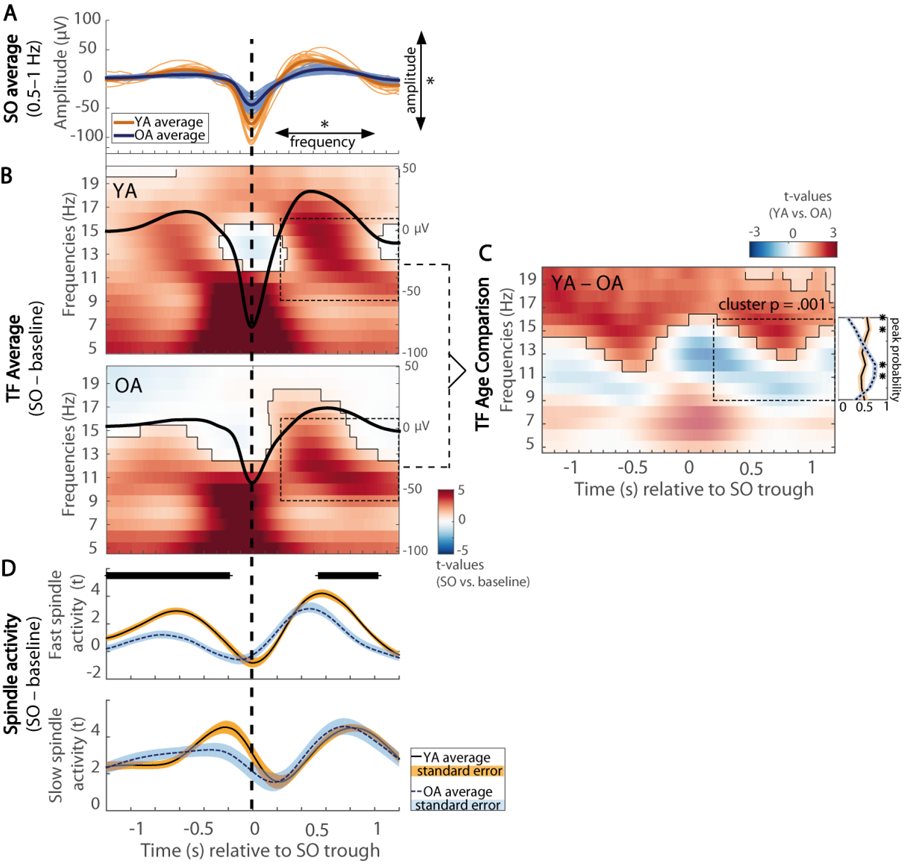
SO–SP coupling is differentially expressed in younger and older adults. (A) Average frontal SOs are depicted for each younger (light orange) and older (light blue) individual. In older adults, the amplitude of SOs is reduced and the frequency is decreased. (B) Differences in wavelet power for SO trials (respective trough ± 1.2 s) compared to baseline trials without SOs are depicted (in *t*-score units, reference window for later analyses outlined by dashed black line). Significant clusters (cluster-based permutation test, cluster α < 0.05) are highlighted and outlined. The average frontal SO for each age group is shown in black (the scale in μV is indicated on the right of each time–frequency graph). In both age groups, EEG activity is modulated as a function of the SO phase. (C) Time–frequency *t*-map illustrating reliable age differences in SO-related power increases compared to baseline trials (cluster *p* < .001). Significant age differences in fast SP power modulation are tied to the SO up-states. The probabilities of frequency peaks (maximal *t*-values) across time (reference window outlined by dashed black line) are shown in the line plot to the right. In older adults the probability of high fast SP activity is significantly reduced during the SO up-state. In contrast, compared to younger adults, a strong increase in slow SP frequencies is more likely (right line plot, all clusters: *p* ≤ .007, marked by asterisks). (D) Temporal evolution of SO-specific power modulations (expressed in *t*-values) in the slow and fast SP range extracted from the time–frequency maps above. In older adults, compared to younger adults, SO-specific fast SP activity is significantly reduced during and following the SO peak (significant clusters marked in black). YA: younger adults; OA: older adults; SO: slow oscillation; SP: spindle; TF: time–frequency.

In younger adults, we observed increases in fast SP power (12.5–16 Hz) during the SO up-state but not during the down-state (Figure 4B). Fast SP power increased most strongly during and shortly after the SO peak following the down-state (400 to 700 ms, SO peak at 480 ms). Although this effect was visible across all chosen derivations (Supplementary Figures 2A and 3A), it was more pronounced over centroparietal sites. Frontal and central slow SP activity (9–12.5 Hz) was increased during the whole SO interval, but peaked during the up- to down-state transition (−500 to 200 ms; Supplementary Figures 2 and 4).

In older adults, as expected, SOs themselves were expressed differently than in younger adults, with a decreased amplitude and frequency (SO amplitude: *z* = 5.69, *p* < .001, *Md*_YA_ = 123.54 μV, *Md*_OA_ = 74.82 μV; SO frequency: *z* = 2.68, *p* = .007, *Md*_YA_ = 0.79 Hz, *Md*_OA_ = 0.78 Hz, Figure 4A). But SO-modulated EEG activity also differed from younger adults (Figure 4B–D). To identify age-specific characteristics of this global pattern we directly compared the time–frequency patterns (i.e., the SO–baseline power differences) of both age groups (Figure 4C). Further, the temporal evolution of frontal slow SP (9–12.5 Hz) and central fast SP power modulation (12.5–16 Hz; averaged *t*-values of the SO–baseline contrast within the respective frequency band) was extracted for both age groups and compared by means of a cluster-based permutation test (Figure 4D). Finally, as studies so far have indicated that SP activity during the SO up-state is critical for memory consolidation^24,28^, we precisely determined in which frequencies SP activity was modulated most strongly during the SO up-state (cf. Methods). We thus compared the probability of frequency peaks between 9 and 16 Hz (maximal *t*-values of the SO–baseline contrast) during the SO up-state (0.2–1.2 s after the SO trough) between younger and older adults (line subplot to the right of Figure 4C).

Over all derivations, power in higher frequencies (16–20 Hz) was globally reduced during SOs in older compared to younger adults whereas fast SP power differences (12.5 – 16 Hz) were reduced during the SO up-states only (all significant positive clusters: *p* ≤ .015, Figure 4C, Supplementary Figure 2). In older adults, the power peak over central sites during the up-state was more stretched in time and shifted to lower frequencies (9–13 Hz, i.e., the slow SP frequency band, Figure 4B). Moreover, SO-specific fast SP activity was reduced during and following the positive SO peak (SO peak at 618 ms; significant cluster: 459 ms – 927 ms; cluster *p* = .013, Figure 4D). In contrast, older adults revealed a strong increase in slow SP activity during the SO up-state (Figure 4B and 3D). Both the age comparison of the global time–frequency pattern (Figure 4D) but also the age comparison of slow SP activity modulated during SOs (Figure 4D, lower panel) indicated a similar SO-specific slow SP power increase in both younger and older adults (all clusters: *p* ≥ .073). Nevertheless, while younger adults showed an almost equal modulation of slow and fast SP power during the SO up-state, older adults displayed a stronger emphasis on lower frequencies: compared to younger adults, fast SP activity was significantly less and slow SP activity significantly more increased during the SO up-state (all clusters: *p* ≤ .007, line subplot to the right of Figure 4C).

To summarize, we found a characteristic SO–SP coupling pattern in younger adults that was marked by a strong increase in fast SPs coupled to the SO peak as well as an increase in slow SPs during the up- to down-state transition. This pattern changed significantly across the adult lifespan. Within this ‘aged’ SO–SP coupling, three characteristics were striking: First, fast SP modulation was reduced overall in the elderly. Second, a reliable fast SP increase comparable to that of younger adults was limited to the time period *before* the SO peak. Together with an overall reduced frequency of SOs, fast SPs were thus no longer precisely tied to the SO peak. Finally, slow SPs were similarly modulated in younger and older adults. Nevertheless, as fast SP power during the up-state was less increased in old age, slow SP power was proportionally more strongly increased in older adults, resulting in a pattern that was characterized by an emphasis on slow SP power at the end of the SO up-state.

### Reduced memory retention is associated with dispersed SO–SP coordination in younger adults

After establishing age-related changes in SO–SP coordination during the SO up-state, we asked how these alterations relate to the ability to retain memories across sleep. As central fast SP activity tied to the SO up-state is critically involved in memory consolidation^24,28^, we focused our analyses on SP frequencies during the SO up-state (9–16 Hz, 0.2–1.2 s) and on electrode Cz, where we observed a lack of SO–fast SP coupling in older adults (Figures 3 and 4; for frontal channels and Pz, see Supplementary Figure 5). To minimize potential biases in SP activity arising from preceding SOs^19^, all analyses were restricted to the up-state following the SO trough.

To quantify memory retention, we measured each participant’s ability to recall scene–word associations that had been successfully encoded the day before. We calculated the percentage of correctly recalled items during delayed recall relative to all items that were correctly retrieved during immediate recall before sleep. Whereas younger and older adults did not differ with regard to their learning performance on Day 1 (t(59) = 1.62, *p* = .11; *M_YA_* = 53.83 %, *SD_YA_* = 20.80 %; *M_OA_* = 45.80 %, *SD_OA_* = 19.02 %), memory consolidation was significantly reduced in older adults. On Day 2, younger adults retained on average 90.58 % of the previously learned pairs *(SD_YA_* = 7.64 %). However, older adults showed significantly worse memory retention across sleep (*M_OA_* = 71.15 %, *SD_OA_* = 13.74 %, t(56) = 6.52, *p* < .001; Figure 5B).

**Figure 5.**
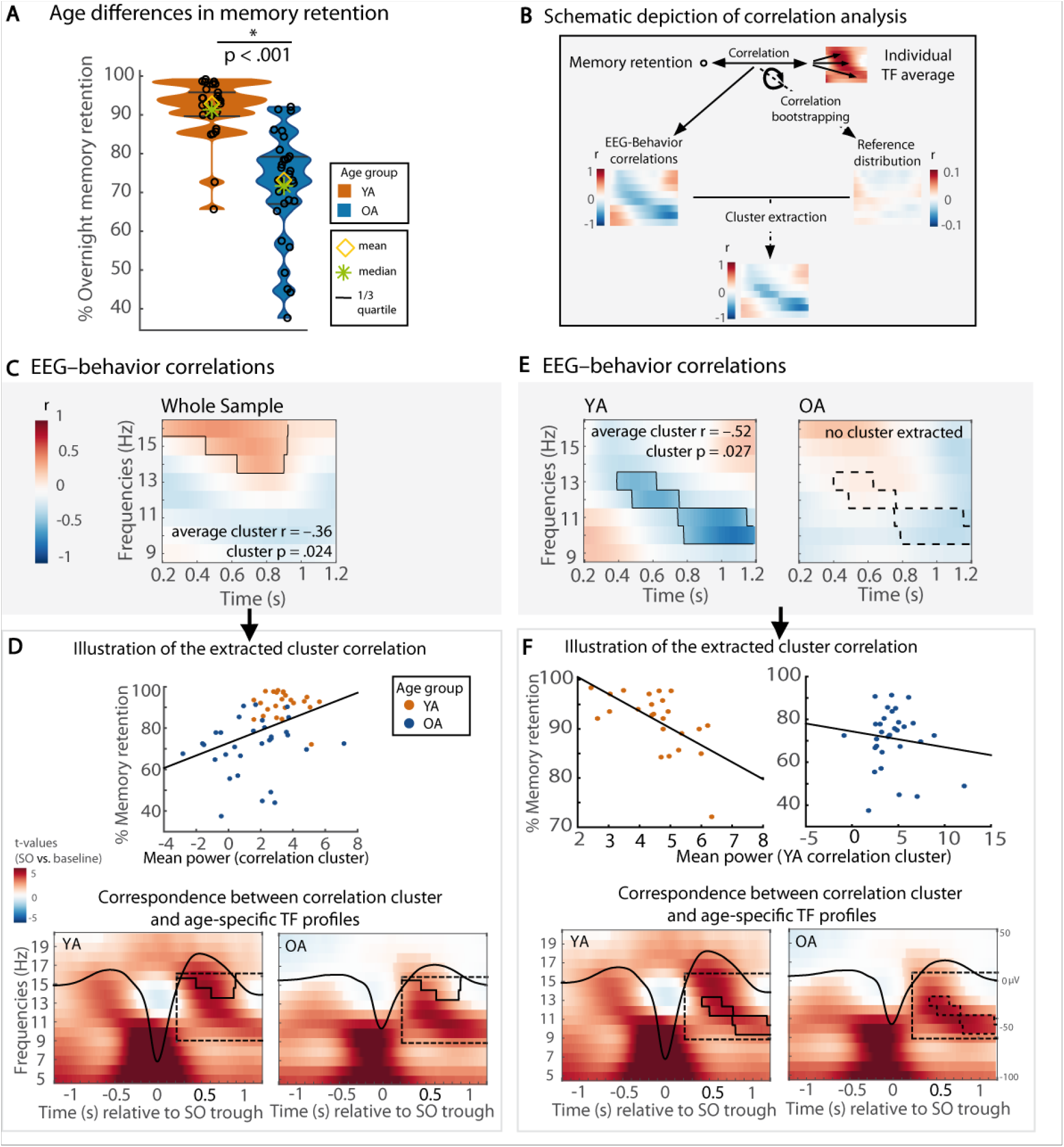
SO–SP coordination relates to memory retention. (A) Age-specific distribution of memory retention in younger (orange) and older adults (blue). Older adults’ overnight memory retention is significantly reduced. (B) Extraction of significant correlation clusters between SO-specific EEG activity and memory retention was achieved by contrasting a test statistic (i.e., the correlation for each time–frequency point) against a reference distribution of bootstrapped EEG–behavior correlations. (C) Correlations between memory retention and neuronal activity during SO up-states are shown for the whole sample (significant positive correlation cluster highlighted and outlined). (D) Upper plot: Scatter plot depicting the positive association between averaged SO-related power modulations (t-values) in the correlation cluster and memory retention (least-squares fit line shown in black). Lower plots: Correspondence between the correlation cluster (outlined in black) and the age-specific time–frequency profiles (reference window for the correlation analyses marked by dashed black line). Fast SP activity during the SO peak, that is typically more expressed in younger adults, was related to better retention of previously learned information. (E) In younger adults, a significant negative correlation cluster (highlighted and outlined) was detected. (F) Upper plots: Scatter plots depicting the negative association between mean power in the correlation cluster and memory retention for both age groups (least-squares fit line shown in black). Lower plots: Correspondence between the correlation cluster and the age-specific time–frequency profiles. The significant cluster found in younger adults corresponds to the power increases specific for older adults. Note. Scatter plots (D and F) only serve illustrative purposes. Hence, no significance is stated in the respective subplots. YA: younger adults; OA: older adults; SO: slow oscillation; SP: spindle; TF: time–frequency.

Correlational analyses between SO–baseline power differences (expressed in *t*-values) and memory retention were performed separately for each time–frequency point. Significant associations were identified by testing against a bootstrapped reference distribution of the EEG–behavior correlations. In this way, we could identify significant correlation clusters that represent the association between memory retention and a specific pattern of EEG activity modulated during the SO up-state (cf. Methods, see Figure 5A for a schematic depiction of the performed correlation technique). Across both age groups, we identified a significant positive correlation cluster (cluster *p* = .024, mean *r* = .36, Figure 5C): more fast SP activity during the SO up-state, which is in general more typical for SO–SP coupling in younger adults, was associated with better memory retention (Figure 5D for an illustration of the observed effect).

To further probe the association between memory retention and SO-modulated EEG activity unbiased by age differences in behavior and/or SO–SP coupling, we then proceeded to conduct the very same analysis within each age group separately. While the positive association between fast SP power during the SO up-state and memory retention did not result in a reliable cluster in the older adults sample (Figure 5E), we found a significant negative correlation cluster in younger adults (cluster *p* = .028, mean *r* = –.52, Figure 5C). Younger adults showing a SO-specific power modulation with less emphasis on fast SP activity during the SO peak but rather on lower frequencies (9–13 Hz) during and following the SO peak showed worse overnight retention of memories (see Figure 5D for an illustration of the negative correlation). Although no significant correlation cluster was extracted in older adults, the average correlation within the cluster identified in younger adults did not differ significantly between age groups *(r_OA_* = –.11, *z* = –1.64, *p* = .10, Figure 5F). This suggests that the same functional association between SO-modulated SP activity and memory retention exists in both younger and older adults. To further illustrate the functional significance of the EEG–behavior association, we checked for the correspondence between the correlation cluster and the age-specific time–frequency profiles. As depicted in the lower panel of Figure 5F, the negatively associated power increases in younger adults overlapped with the typical SO-related activity pattern of older adults that is marked by a shift to lower SP frequencies and a more stretched power peak sustained until the end of the SO up-state. Younger adults who did *not* display an ‘age-typical’ coupling pattern tended to be worse at retaining previously learned information overnight. As a control, we ran the same analysis for a second time window (–500 to 200 ms) during the SO up- to down-state transition that reflects SO-modulated slow SP activity. Here, no significant associations with memory retention were observed after controlling for multiple comparisons (cluster *p* = .169, Supplementary Figure 6).

To conclude, we found evidence for an association between the coordination of SOs and SPs and memory retention. In general, high levels of fast SP power during and following the positive SO peak were related to better memory retention. Moreover, our results suggest that inter-individual differences in SO–SP coupling as maximized *between* age groups also drive differences in memory retention *within* age groups. Dispersed SO–SP coupling, which was characteristic of older adults, already predicted worse memory retention among younger adults.

### Structural brain integrity in old age promotes ‘youth-like’ SO–SP coupling

In line with previous reports on the influence of age-related brain atrophy on the generation of sleep oscillations^35,36^, we finally asked whether the identified age-specific modulation of SP activity during the SO up-state is associated with measures of brain volume in specific regions of interest (ROI). As ROIs we included brain regions that are considered source regions for the generation of SOs and SPs^9,11^ and proposed to be involved in memory processing during sleep^47^. We also included an occipital control region (Figure 6A). To exclude any age-related confounds, measures of brain volume were derived by using voxel-based morphometry (VBM) and correcting the extracted ROI-specific measures for total intracranial volume (TIV)^48^. Again, we focused our analysis on frequencies between 9 and 16 Hz and a time window between 0.2 and 1.2 s after the SO trough. Whereas the cluster-corrected correlation analysis (cf. Figure 5A, Methods) did not indicate any significant association between brain volume and EEG power in younger adults (all clusters: *p* ≥ .073; Supplementary Figure 8), we observed a very specific pattern in older adults.

**Figure 6.**
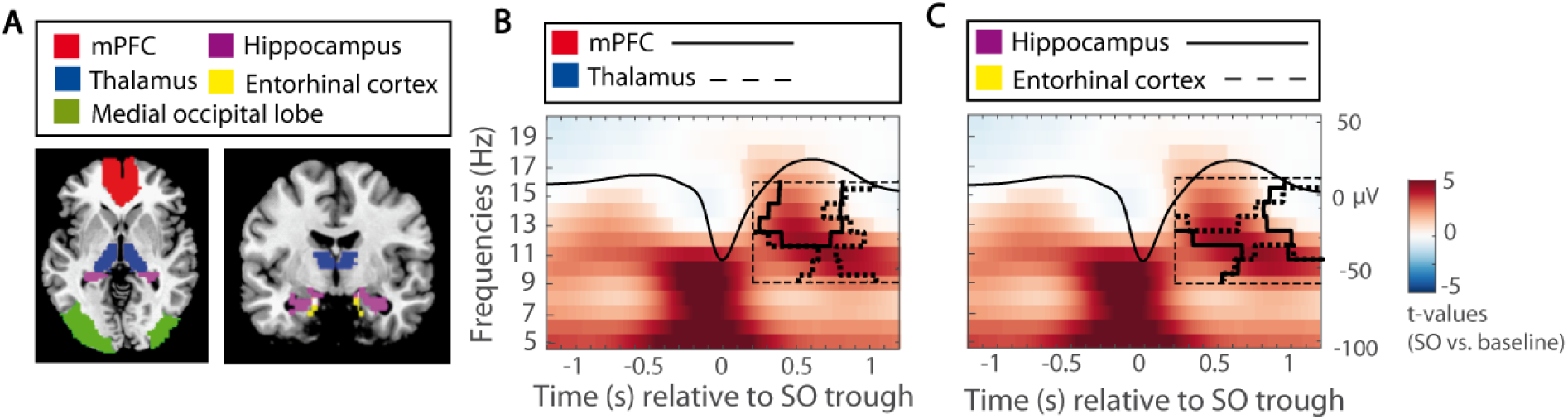
Structural brain integrity in old age promotes ‘youth-like’ SO–SP coupling. (A) VBM measures are extracted from ROIs to obtain individual values of brain volume (ROI masks overlaid in color). (B) In older adults, greater volume in the mPFC and the thalamus is associated with increased fast SP activity at the peak of the SO. Significant positive correlation clusters (controlled for multiple comparison using a cluster-corrected correlation approach, cluster α = 0.05) are outlined by solid (mPFC) and dashed black lines (thalamus). Underlying time–frequency profiles are only plotted for illustration (in *t*-score units, reference window for the correlation analysis outlined by dashed black line). (C) Greater volume in the hippocampus (black solid line) and entorhinal cortex (black dashed line) is positively correlated with more global increases in EEG activity during the SO up-state. mPFC: medial prefrontal Cortex; VBM: voxel-based morphometry; ROI: region of interest; SO: slow oscillation; SP: spindle.

In older adults, larger medial prefrontal cortex (mPFC) was associated with higher power in the fast SP range during the SO peak (mean *r*_mPFC_ = .43, cluster *p*_mPFC_ = .018). Thalamus volume showed a similar positive relation with spindle activity during the SO peak. This effect, though, was less precise with regard to the frequency range (mean *r*_thalamus_ = .45, cluster *p*_thalamus_ = .013, see Figure 6). For hippocampus and entorhinal cortex even broader effects were observed. Greater volume in these regions was not only associated with higher fast SP power during the SO peak, but generally with more global neuronal activation during the SO up-state. In particular, the observed effect extended to slow SP frequencies at the end of the SO up-state (mean *r*_hippocampus_ = .47, cluster *p*_hippocampus_ = .002; both mean *r*_entorhinal cortex_ = .46, cluster *p*_entorhinal cortex_ = .013 and .031). As expected, no effect was detected when looking at medial occipital lobe volume (all clusters: *p* ≥ .169).

To summarize, older adults with less age-related decline in mPFC and thalamus volume showed a more precise coupling between fast SPs and SOs time-locked to the SO peak – precisely the pattern observed in younger adults. Less age-related decline in medial temporal lobe areas was moreover associated with overall enhanced neuronal activity during the SO up-state. We thus conclude that structural brain integrity in old age promotes ‘youth-like’ SO–SP coupling.

## Discussion

This study investigated whether the coupling of SOs and SPs during sleep changes across the adult lifespan and how these alterations relate to the ability to retain associative memories across sleep. We demonstrate that older adults do not display the central fast SP power increase time-locked to the SO peak that we observed in younger adults. Instead, they exhibit overall reduced fast SP power modulations and a fast SP peak that is shifted before the SO peak. Moreover, SO-modulated SP activity is characterized by a strong emphasis on lower frequencies at the end of the SO up-state. We find evidence that this ‘aged’ coupling pattern coincides with worse memory consolidation, whereas a ‘youth-like’ precision of SO–SP coupling promotes memory consolidation across the entire adult lifespan. Moreover, greater structural integrity in brain regions involved in the generation of SOs and SPs relates to an intact ‘youth-like’ SO–SP coupling in old age. Overall, our results resonate well with a recent report on age-related differences in SO–SP phase–amplitude couplings and their functional significance for memory retention^39^. We extend these findings by applying analysis methods that are sensitive to age-related shifts in both the time and frequency domain and explicitly base our analyses on the differentiation between slow and fast SPs. Together with Helfrich and co-authors^39^, we are the first to demonstrate that an age-related dispersion of SO–SP coupling, as caused by structural brain atrophy, relates to impaired memory consolidation in old age.

### Age-related dispersion of SO–SP coupling

Effective neural communication during NREM sleep is governed by SOs that, during their up-states, initiate the generation of thalamic fast SPs^11^. Here, we found that a generally constant pattern of SP–SO co-occurrence exists across adulthood, although SO and SPs did not co-occur in a one-to-one fashion (Figure 2). Our results imply that: (1) Although it appears that thalamic SP generation is triggered by frontal SOs^11,12^, at least one other mechanism initiating SPs exists. (2) SO-induced SP initiation is not a necessity, but rather a rare occurrence. Nevertheless, if SOs and SPs co-occur, they do so in a precisely timed manner, as discussed next.

Consistent with a wide range of studies, we identified a precisely timed hierarchical structure of SOs and SPs in younger adults with fast SPs appearing during the highly depolarized SO peak and slow SPs occurring at the up-to down-state transition^16,19,20,23^. SOs themselves change their appearance in old age and exhibit reduced amplitudes and slower frequencies in older adults (Figure 4A)^31,33^. Flatter SOs have previously been associated with less synchronous neuronal switches between phases of depolarization and hyperpolarization^34^. As these phases precisely time the generation of SP events themselves, SO–SP coordination should, as a consequence, disperse in old age.

Indeed, consistent with recent reports^39,44^, in older adults EEG activity during SOs was differentially modulated than in younger adults. (1) In line with previously reported reductions in central fast SPs in old age^32,49^, fast SP increases during the SO up-state proved less pronounced in older adults. (2) If fast SPs appeared, they were no longer precisely tied to the SO peak as observed in younger adults. Rather, they occurred before the up-state reached its maximum. (3) Over central sites modulation of EEG activity during the SO up-state dispersed, the power peak was stretched in time and shifted to lower frequencies (9–13 Hz). All things considered, we argue that even in older adults a very specific pattern of SO–SP coupling exists, although its appearance changes compared to younger adults, with more emphasis on lower frequencies occurring later during the SO up-state. Possible causes and mechanisms of this ‘aged’ pattern will be discussed next.

### Structural correlates of SO–SP coupling

The process of SO–SP coupling involves intact communication between different brain regions^11^. SOs, generated mainly in prefrontal areas, propagate throughout the brain^9^ and potentially trigger the initiation of fast SPs in the thalamus^11^. Here, we indicate that in old age structural integrity in the mPFC accounts for a ‘youth-like’ SO–SP coupling (Figure 6). This is in line with Helfrich and colleagues (2018) who recently demonstrated that greater mPFC volume relates to a more precise coupling of fast SPs to the SO peak^39^. Expanding on that observation, we show that additional brain regions are associated with inter-individual differences in SO–SP coupling. Greater thalamus volume, the main source region of fast SP generation^13^, related to more SP activity during the SO peak. The specificity of these effects strongly suggests that structural integrity in both mPFC and thalamus is crucial for pacing a ‘youth-like’ SO–SP succession. In contrast, hippocampus and entorhinal cortex volumes were linked to both fast SP activity coupled to the SO peak and slow SP activity occurring at the up-to down-state transition. Both hippocampus and entorhinal cortex are key players in the overnight consolidation of memories^2^. In particular, sharp-wave ripples are generated within the hippocampus and thereby complete SO–SP coupling^16^ as the basis for successful memory reprocessing during sleep^25,26^. Thus, we speculate that the association between structural integrity of the medial temporal lobe and a successful coordination of oscillatory processes during sleep is indicative for sharp-wave ripple activity.

In our analyses, significant associations between SO–SP coupling and VBM measures were only found within the older sample. Importantly, this does *not* imply that those brain areas do not contribute to an effective hierarchical nesting of neuronal oscillations during sleep in younger adults. We rather suggest that brain integrity in mPFC, thalamus and medial temporal lobe areas forms the basis for an effective ‘youth-like’ SO–SP coupling independent of age. Under conditions of impaired structural integrity in these regions, as is the case in aging^48^, the precise timing of SOs and SPs vanishes. This reasoning is in line with Nyberg and colleagues (2012) who argue that the individuals who experience fewer age-related structural brain changes are more likely to show neural activity patterns that resemble those of younger adults and that are associated with better performance^50^. The functional implications of an ‘aged’ SO–SP coupling will be discussed next.

### Functional significance of precise SO–SP coupling

Current theory holds that hippocampal–neocortical communication during sleep is only enabled by the precise hierarchical coordination of SOs, SPs, and hippocampal sharp-wave ripples^51,52^. This is achieved by locking fast SPs to the SO peak and further nesting sharp-wave ripples within the troughs of fast SPs^16^. Thus, memories initially dependent on a temporary store in the hippocampus become more reliant on neocortical areas, where they are permanently stored^4,22,53^.

Here, we show that, across younger and older adults, ‘youth-like’ SO–SP coupling tended to coincide with better memory retention (Figure 5C and 5D). In contrast, ‘aged’ SO–SP coupling was associated with worse memory retention in younger adults (Figure 5E and 5F). Our results support previous evidence showing that the precisely timed interaction of neural rhythms indeed supports sleep-dependent memory consolidation^25–28,39,44^. Brain activity is globally synchronized during the SO up-state and maximized during the SO peak^14^. This constitutes a moment of effective interregional brain communication. Fast SPs occurring at this very moment induce a massive influx of Ca^2^+ into excited neurons. This enables synaptic plasticity and promotes the stabilization and integration of memory representations^54^. As most recently demonstrated, fast SPs are misaligned in older adults, resulting in worse memory retention^39^. But what is it that makes the ‘aged’ SO–SP coupling detrimental?

First, the temporal dispersion of ‘aged’ SO–SP coordination in older adults is striking. In older adults, the fast SP power increase during the SO up-state reaches its maximum *before* the SO peak (Figures 3 and 4). This may be explained in two ways: (1) During flatter SOs, as observed in older adults, the switch between up- and down-states is more variable^31,33,34^ and as a result SP initiation is also less precise. (2) The mechanism initiating SPs might be similarly timed in younger and older adults, but as SOs are slower in older adults, fast SPs appear too early during the SO up-state. Here, global brain synchrony has not reached its maximum yet and interregional brain communication might not be optimal for reorganization of memories on a brain system level.

The second characteristic of ‘aged’ SO–SP coupling is a power shift to lower frequencies in the slow SP band at the end of the SO up-state (Figure 3). At young age already, this shift related to worse memory retention (Figure 5, Supplementary Figure 7). Recent evidence suggests that fast and slow SPs are generated via distinct mechanisms^55,56^ and have different functional significances. While the occurrence of fast SPs probably reflects thalamo–cortical information processing necessary for memory transfer, slow SPs may mirror cortico-cortical communication^57^. Slow SPs per se may not be sufficient to enable effective memory consolidation during sleep^2,19,21,28^. When focusing our analysis on the up-to down-state transition, where slow SPs preferentially occur^19,20^, no association with memory retention was detected (Supplementary Figure 6). To conclude, our results replicate the finding that a precise ‘youth-like’ SO-fast SP coupling is beneficial for memory consolidation^39,44^. Vice versa, ‘aged’ coupling coincides with worse memory consolidation already in young adulthood. This lines up with the concept of “brain maintenance”^50^ stating that cognitive functioning is not only determined by maintained structural integrity across the adult lifespan, but also by the preservation of functionally specific processes and networks^50,58,59^. Maintenance of ‘youth-like’ processing during both encoding^58^ and memory retrieval^59^ has previously shown to relate to better long-term memory in older adults. Here we complement this view by demonstrating that consolidation processes also benefit from maintained brain mechanisms during sleep.

## Methods

### Participants and procedure

#### Participants

Thirty-four healthy younger adults (19–28 years) and 41 healthy older adults (63-74 years) participated in the experiment. Due to technical failures, data collection from 4 younger and 4 older adults could not be completed. Accordingly, the final sample for our behavioral analysis consisted of 30 younger (*M_age_* = 23.7 years, *SD_age_* = 2.6; 17 females) and 37 older adults (*M_age_* = 68.92 years, *SD_age_* = 3.04; 16 females). For some participants parts of the neural data (PSG or MRI) were missing or of bad quality. Final PSG analyses were hence conducted with 24 younger (*M_age_* = 23.61 years, *SD_age_* = 2.55; 13 females) and 31 older adults (*M_age_* = 68.63 years, *SD_age_* = 3.10; 15 females), VBM analyses with 24 younger (*M_age_* = 23.61 years, *SD_age_* = 2.55; 13 females) and 29 older adults (*M_age_* = 68.64 years, *SD_age_* = 3.10; 14 females). The different samples did not differ with regard to any behavioral measure.

All participants were right-handed native German speakers with no reported history of psychiatric or neurological disease, or any use of psychoactive medication. To screen for cognitive impairments in the sample of the elderly, all older adults completed the Mini-Mental State Examination (MMSE; *M* = 29.24, *SD* = 1.12, *Range:* 26–30)^60^ and passed a brief memory screening beforehand. General subjective sleep quality was controlled by assessing the Pittsburgh Sleep Quality Index (PSQI)^61^ and did not show differences between the two age groups. The study was approved by the Ethics Committee of the *Deutsche Gesellschaft für Psychologie* (DGPs) and conducted at the *Max Planck Institute for Human Development* in Berlin. After being informed about the complete study procedure, all participants gave written consent to their participation in the experiment.

#### Experimental Procedures

The present data were derived from a series of studies investigating age-related differences in the encoding, consolidation, and retrieval of associative memories (see Fandakova et al., 2018, for the effects of age and memory quality on false memory retrieval^62^). Core of the experimental design was a paired-associative scene–word memory paradigm, consisting of a learning session on the first day (Day 1) as well as a delayed cued-recall task approximately 24 hours later (Day 2) (see Figure 1, for illustration of the study procedure). During the nights before and after learning (experimental nights PRE and POST) sleep was monitored at participants’ home using ambulatory PSG. Prior to the first experimental night an adaptation night familiarized the participants with the PSG procedure. Structural MRI data were collected on Day 2. Furthermore, EEG was recorded during learning on Day 1. Additionally, functional MRI data were collected during delayed recall on Day 2. Neither the EEG nor the fMRI data are included in the present report. Participants completed a short cognitive screening prior to participation in the main study, but did not engage in any other behavioral task during the course of the experimental procedure described above.

#### Day 1 – Encoding and immediate cued-recall

During initial study, scene–word pairs were presented on a black background for 4000 ms. Scene–word pairs were randomized combinations of indoor and outdoor scenes and concrete nouns. Participants were instructed to remember the scene–word pair using a previously trained imagery strategy and to indicate how well they were able to form an integrated image of the scene–word pair. Cued-recall blocks followed immediately after the initial learning phase. Scenes served as cues for participants to verbally recall the associated word. Recall time was not constrained. Independent of recall accuracy, the correct scene–word pair was presented again for 3 seconds and participants were instructed to use this opportunity to remember the word–scene pair by forming an integrated image of the two. Finally, participants completed a final cued-recall test without feedback at the end of Day 1. This phase served as a measure of learning performance. Importantly, task difficulty was adjusted between the age groups to achieve comparable recall success of approximately 50 % in each age group. This was done in two ways. First, younger adults learned 440 pairs, whereas older adults learned 280 pairs on Day 1. Second, younger adults completed one cued-recall block with feedback, whereas older adults completed two cued-recall blocks with feedback. Further details about the encoding task and the stimulus set are described by Fandakova et al. (2018)^62^.

#### Day 2 – Delayed cued-recall

Delayed cued-recall of the scene–word pairs consisted of four blocks of 70 experimental trials each and was conducted inside a 3T MRI scanner. For the older age group, all of the 280 studied pairs were presented. For younger adults items were chosen with regard to their learning history. This resulted in a selection of pairs, half of which had been recalled in the criterion cued-recall the day before. The 280 pictures were displayed for 3500 ms. Meanwhile participants had to decide via keypress if they still remembered the corresponding word (“remembered” vs. “forgotten”). Afterwards, four letters were presented for 3500 ms. One of these was the second letter of the correct word and had to be selected by pressing the corresponding key. If the according word had been forgotten or none of the letters fitted their remembered answer word, participants were told to choose one of the letters at random. Answers during delayed recall were only classified as correct if participants both responded that they remembered the word *and* selected the right letter afterwards.

### Sleep EEG data acquisition and analyses

#### Data acquisition

During the two experimental nights before (PRE) and after learning (POST), sleep was recorded using an ambulatory PSG device (SOMNOscreen plus; SOMNOmedics, Germany). Eight scalp electrodes were attached according to the international 10–20 system for electrode positioning (Fz, F3, F4, C3, C4, Cz, Pz, Oz)^63,64^ along with two electrodes on the mastoids A1 and A2 that later served as the offline reference. All impedances were kept below 6 kΩ. Data were recorded using Cz as the online reference for all EEG derivations and AFz as ground. Additionally, a bilateral electrooculogram (EOG) was assessed. Two submental electromyogram channels (EMG) were positioned left and right inferior of the labial angle and referenced against one chin electrode. Electrical activity of the heart was recorded using two electrocardiogram channels (ECG). All electrodes were attached to a small recorder fixed with two straps either to the front or back of the torso depending on the participant’s sleep habits. EEG channels were recorded between 0.2–75 Hz with a sampling rate of 128 Hz.

#### EEG pre-processing

Data was preprocessed using *BrainVision Analyzer 2.1* (Brain Products, Germany). Here, all EEG channels were re-referenced against the average of A1 and A2. Afterwards, sleep stages 1 and 2, slow-wave sleep, REM sleep, awakenings, and body movements were visually scored in 30-second epochs according to the standard criteria suggested by the American Academy for Sleep Medicine (AASM)^45^. Participants provided markers for switching the light off and on again, narrowing the time period to be analyzed. Visual determination of sleep onset and offset took place when participants did not remember to set the markers (*n* = 5). Total sleep time (TST) was calculated as time spent in stage 1, 2, SWS, and REM sleep. Wake after sleep onset (WASO) was defined as the time participants were awake between sleep onset and final morning awakening.

Further analyses of the sleep EEG data were conducted using *Matlab* R2014b (Mathworks Inc., Sherbom, MA) and the open-source toolbox *Fieldtrip*^65^. Bad EEG channels were visually rejected. For the remaining channels, artefact detection was implemented on 1-second long segments. Segments that were visually identified as body movement or exhibited amplitude differences of more than 500 μV were marked as bad. To further exclude segments that strongly deviated from the observed overall amplitude distribution, mean amplitude differences for each segment were z-standardized within each channel. Segments with a z-score of more than 5 in any of the channels were also excluded.

#### Spindle detection

SPs were detected using an established automated algorithm^19,20,66^. For all NREM epochs (i.e., stage 2, and slow-wave sleep), EEG data were band-pass filtered using a 6th-order Butterworth filter (in forward and backward direction to prevent phase distortions) between 9 and 12.5 Hz for the detection of slow SPs, respectively 12.5 and 16 Hz for fast SPs^67,68^. The root-mean-square (RMS) representation of the signal was calculated using a sliding window of 200 ms at every sample point. Afterwards, the RMS signal was smoothed by applying a moving average of 200 ms. To increase sensitivity and specificity of our algorithm^69^, we accounted for individual differences in EEG amplitude by anchoring the SP identification on individually determined amplitude thresholds. A potential SP was tagged if the amplitude of the smoothed RMS signal exceeded its mean by 1.5 SD of the filtered signal for 0.5 to 3 seconds. SPs with boundaries closer than 0.25 seconds were eventually merged in the following way: in each processing run, starting with the smallest boundary difference, two SPs were merged if the resulting SP event remained within the time limit of 3 seconds. As soon as a SP was combined with another one, it could not be merged with another SP in the very same run anymore. Only if all boundary differences were revised and no further merging of putative SPs was possible, the new resulting SP events were processed in a next run. Starting again with the smallest boundary difference, two putative SPs could be merged if the merged event remained within the required time limit. These runs were repeated until no further merging was possible. Finally, only SPs not overlapping with previously marked artefact segments were considered SPs.

#### Slow oscillation detection

Detection of SOs at frontal electrodes was based on Mölle et al. (2002)^23^ and Ngo et al. (2013)^70^. For all NREM epochs, EEG data was band-pass filtered between 0.2 and 4 Hz using a 6th-order Butterworth filter (in forward and backward direction). The whole signal was then divided into negative and positive half-waves that were separated by zero-crossings of the filtered signal. The combination of a negative half wave with the succeeding positive half wave was considered a putative SO when its frequency was between 0.5 and 1 Hz. The amplitude of each potential slow wave was calculated as the distance between the SO trough and positive peak, defined as its maximal negative and positive potential, respectively. As in the SP detection, adaptive amplitude thresholds were defined separately for each participant: Putative SOs exceeding a trough of 1.25 times the mean trough of all putative SOs as well as an amplitude of 1.25 times the average amplitude of all potential SOs, were marked. Only artefact-free SOs were considered in further analyses.

#### Statistical analysis

Not all sleep variables used in our analysis followed a normal distribution. To identify age differences in the sleep architecture and the expressed sleep oscillations between younger and older adults, non-parametric Mann-Whitney *U* Tests for independent samples were calculated and median and quartile values of the variables were reported.

##### Temporal relation between detected SO and SP events

The general temporal relation between SO and SP events was calculated by determining the proportion of SPs whose center occurred in an interval of ± 1.2 s around the trough of the identified SOs, and the amount of SOs with SPs whose center occurred within ± 1.2 s around the respective trough of the oscillation. The time window of ± 1.2 s was chosen to cover one whole SO cycle (0.5–1 Hz, i.e., 1–2 s). The exact timing of SO and SP events was visualized by peri-event time histograms (PETHs) of fast and slow SP centers (seed events) occurring within a time interval of ± 1.2 s around each SO trough (target event). Probabilities of seed event occurrence were summed within bins of 100 ms and normalized to add up to 100 %. Following the same pipeline, PETHs were also calculated for intervals of ± 5 s to demonstrate the general specificity of SP appearance to the actual presence of SOs (Supplementary Figure 3 and 4). To test for the temporal stability of the PETHs, we implemented a randomization procedure by randomly shuffling the temporal order of the PETH bins 1000 times. The resulting surrogates were averaged for each individual and tested against the original PETHs across subjects using dependent sample *t*-tests. Control for multiple comparisons was achieved by applying a cluster-based permutation test with 5000 permutations^71^.

##### Time–frequency analyses

To describe the temporal association between SPs and SOs, we selected 6-second long artefact-free trials from the EEG data, centered on the trough of the SOs ± 3 s. The longer time segments were selected to prevent filter artefacts during later analysis steps (e.g., due to wavelet filtering). All analyses described in the following were first conducted separately for SOs detected at the two frontal derivations F3 and F4, and then averaged. time–frequency analyses were performed at frontal electrodes as well as at electrodes Cz and Pz. To achieve an appropriate baseline contrast allowing for the interpretation of SO specific power decreases and increases, we matched every detected SO with a randomly chosen artefact–free time segment of 6 seconds during the same sleep stage as the respective SO. To obtain time–frequency representations of trials with and without SOs, we applied a Morlet-wavelet transformation (12 cycles) to the unfiltered EEG data of SO and baseline trials between 5 Hz and 20 Hz in steps of 1 Hz and 2 ms. Trials with and without SOs were then contrasted for each subject using independent-sample *t*-tests. The resulting *t*-maps reflect the increase/decrease in both the fast and slow SP frequency range for trials with SOs, compared to trials without. Due to the high prevalence of SOs during slow-wave sleep, the number of identified baseline trials sometimes differed from the number of SO trials during this sleep stage. Thus, to calculate the within-subject contrasts, we drew 100 random sets of baseline and SO trials while maintaining the ratio of stage 2 to slow-wave sleep trials (average trial number: 599 ± 225). *t*-maps for all these random trial sets were averaged for each subject. These *t*-maps were further averaged across F3 and F4 SOs for each participant before subjecting the results to group-level analyses. Separately for each age group, *t*-maps were tested against zero using a cluster-based permutation test with 5000 permutations in a time window of –1.2 to 1.2 s^71^.

##### Age comparison of SO–SP coupling

Age differences in SO-modulated SP activity were quantified in three ways: (1) The time–frequency *t*-maps (SO vs. baseline trials) of both age groups were compared using a cluster-based permutation test with 5000 permutations. (2) The time course of frontal slow SP (9–12.5 Hz) and central fast SP power modulation (12.5–16 Hz) was extracted by averaging the *t*-values of the SO–baseline contrast within the respective frequency band. SO-specific SP power modulation was extracted at each time point during the SO (SO trough ± 1.2 s) for both age groups and compared by means of a cluster-based permutation test with 5000 permutations. (3) Finally, we determined the frequencies with strongest power modulation during the SO up-state. We focused on a time–frequency window covering the whole SP range (9–16 Hz) and the entire SO upstate (0.2–1.2. s). For each frequency, we determined the peak power modulation (i.e., the maximal *t*-values of the SO–baseline contrast) during the SO up-state. For each subject, the maximum *t*-values across frequencies were scaled between 0 (lowest *t*-value) and 1 (peak modulation). The resulting distribution reflects the probability of observing the maximum power differences between SO and baseline trials at a given frequency during the SO up-state. Age differences in the peak probability distribution were calculated using a cluster-based permutation test with 5000 permutations.

##### EEG-behavior correlation

As we aimed to identify the functional significance of the detected SO–SP association during the SO up-state, we then correlated each participant’s individual time–frequency *t*-maps for the comparison with/without SOs with the participant’s ability to retain previously learned information overnight (cf. Figure 5A for a schematic depiction of the correlation analysis). For each time–frequency point (9–16 Hz, 0.2–1.2. s) a Pearson’s correlation with the behavioral outcome variable was computed. Control for multiple comparisons was achieved using a cluster-corrected correlation approach with 5000 bootstrap samples to create the reference distribution. Correlation analyses were performed across all subjects and within each age group separately.

### MRI data acquisition and structural MRI analyses

Whole-brain MRI data was acquired with a Siemens Magnetom 3T TimTrio machine. For each participant a high-resolution T1-weighted MPRAGE sequence (TR = 2500 ms, TE = 4.77 ms, FOV = 256 mm, voxel size = 1 × 1 × 1 mm^3^) was collected. To get estimates of brain volume in regions of interest (ROI) we conducted voxel-based morphometry (VBM) using statistical parametric mapping software (SPM12, http://www.fil.ion.ucl.ac.uk/spm) and the Computational Anatomy Toolbox (CAT 12, http://www.neuro.uni-jena.de/cat). Images were normalized to Montreal Neurological Institute (MNI) space and segmented into gray matter, white matter, and cerebrospinal fluid. Data were then modulated, i.e., corrected by the volume changes due to spatial normalization. For this, each voxel value was multiplied with the Jacobian determinant derived from the spatial normalization step. Afterwards images were smoothed with a 8 mm full width at half maximum (FWHM) kernel. Total intracranial volume (TIV) was estimated by summing volume of gray matter, white matter, and cerebrospinal fluid. Due to their involvement in sleep-dependent memory processes and especially in the generation of SPs and SOs^5,9,11^, we selected bilateral medial prefrontal cortex (mPFC), thalamus, entorhinal cortex, hippocampus, and, as a control region, the medial occipital lobe (Figure 7A). The mPFC mask was kindly provided by Bryce A. Mander^35^. All other ROIs were defined using the WFU PickAtlas toolbox (http://fmri.wfubmc.edu/software/pickatlas). Measures of gray matter volume in all ROIs were extracted using the REX toolbox (http://web.mit.edu/swg/rex/rex.pdf) and adjusted for differences in TIV based on the formula provided by Raz et al. (2005) (adjusted volume = raw volume – b x (TIV-mean TIV), where b is the slope of the regression of ROI volume on TIV)^48^.

#### EEG-brain structure correlation

The association between structural brain integrity and SP activity coupled to the SO up-state was similar to the cluster-corrected correlation analysis described above (cf. Figure 5A). Each participant’s individual time–frequency *t*-map was related to gray matter volumes in the different ROIs and referenced to a bootstrapped reference distribution (5000 bootstrap samples).

## Code availability

The custom code used for these analyses is available upon reasonable request from the corresponding authors.

## Data availability

The data that our results are based on are available upon reasonable request from the corresponding authors.

## Acknowledgements

This study was conducted within the *‘Cognitive and Neuronal Dynamics of Memory across the Lifespan (ConMem)*’ project at the Center for Lifespan Psychology, Max Planck Institute for Human Development. The research was partially financed by the Max Planck Society. Beate E. Muehlroth was supported by the Max Planck International Research Network on Aging. Markus Werkle-Bergner’s work was supported by a grant from the German Research Foundation (DFG, WE 4269/3-1, Yee Lee Shing as Co-PI) as well as an *Early Career Research Fellowship 2017–2019* awarded by the Jacobs Foundation. Yee Lee Shing and Myriam C. Sander were each supported via Minerva Research Groups awarded by the Max Planck Society. We thank Maren J. Cordi for helping us to set up the technical equipment, Xenia Grande for organizing data collection, Kristina Günther for help in participant recruitment, Julia Delius for editorial assistance, and all student assistants of the *ConMem* project collecting the data. We are grateful to all members of the *ConMem* project for helpful feedback on the analysis. Finally, we thank all study participants for their time.

## Author Contribution

MCS, YF, THG, YLS, and MWB designed the study; BEM performed the experiments; BEM and MWB analyzed the data; BEM and MWB wrote the manuscript; BR and MCS gave conceptual advice. All authors revised the manuscript.

